# Ciprofloxacin facilitates the transfer of XDR plasmids from commensal *E. coli* into epidemic fluoroquinolone-resistant *Shigella sonnei*

**DOI:** 10.1101/767251

**Authors:** Pham Thanh Duy, To Nguyen Thi Nguyen, Duong Vu Thuy, Hao Chung The, Felicity Alcock, Christine Boinett, Ho Ngoc Dan Thanh, Ha Thanh Tuyen, Guy E. Thwaites, Maia A Rabaa, Stephen Baker

## Abstract

The global dissemination of a ciprofloxacin-resistant (cipR) *S. sonnei* clone outlines the mobility of this important agent of diarrheal disease, and threatens the utility of ciprofloxacin as a first-line antimicrobial for shigellosis. Here, we aimed to track the emergence of cipR *S. sonnei* in Vietnam to understand how novel antimicrobial resistant (AMR) *Shigella* clones become established in new locations. From 2014 to 2016, we isolated and genome sequenced 79 *S. sonnei* from children hospitalized with dysenteric diarrhea in southern Vietnam. The novel cipR *S. sonnei* clone displaced the resident ciprofloxacin-susceptible lineage while acquiring resistance against third-generation cephalosporins, macrolides, and aminoglycosides. This process was not the result of a single clonal expansion, as we identified at least thirteen independent acquisitions of ESBL-encoding plasmids. The frequency and diversity of the variable AMR repertoire in an expanding clonal background of *S. sonnei* is unprecedented and we speculated that it was facilitated by horizontal gene transfer from commensal organisms in the human gut. Consequently, we characterized non-*Shigella* Enterobacteriaceae from *Shigella-*infected and healthy children by shotgun metagenomics. We identified a wide array of AMR genes and plasmids in the commensal Enterobacteriaceae, including an *E. coli* isolated from a *Shigella*-infected child with an identical ESBL plasmid to that characterized in the infecting *S. sonnei*. We confirmed that these AMR plasmids could be exchanged between commensal *E. coli* and *S. sonnei* and found that supplementation of ciprofloxacin into the conjugation media significantly increased the conjugation frequency of IncI/*bla*_CTX-M-15_, IncB/O/*bla*_CTX-M-27_ and IncF/*bla*_CTX-M-27_ plasmids. In a setting with high antimicrobial use and a high prevalence of AMR commensals, cipR *S. sonnei* may be propelled towards pan-resistance by adherence to outdated international treatment guidelines. Our work highlights the role of the gut microbiota in transferring resistance plasmids into enteric pathogens and provides essential data to restrict the use of ciprofloxacin globally.

## Introduction

*Shigella* is one of the leading bacterial agents of diarrhea globally; responsible for >165 million diarrheal episodes annually, shigellosis principally affects children in low- to middle-income countries (LMICs) ^1^. Correspondingly, it has been estimated that shigellosis is responsible for 28,000-48,000 deaths in children aged <5 years annually ^2^. In 2014, the Global Enteric Multicenter Study (GEMS) identified *Shigella* as one of the top four pathogens associated with moderate-to-severe diarrheal disease in young children in sub-Saharan Africa and South Asia ^3^.

The genus *Shigella* comprises of four species: *S. flexneri, S. sonnei, S. dysenteriae*, and *S. boydii*, but the current international *Shigella* landscape is dominated by *S. flexneri* and *S. sonnei*. Of these two species, *S. sonnei* is increasingly being isolated, replacing *S. flexneri* as the predominant *Shigella* species in many LMICs in Asia, Latin America, and the Middle East ^4^. Unlike other *Shigella, S. sonnei* exists as a single serotype and has a population structure encompassing five lineages, of which lineage III successfully disseminated globally from the 1970s onwards. A key event facilitating the success of this lineage was the acquisition of a multi-drug resistance (MDR) phenotype, which distinguishes this population ^5,6^. Antimicrobials are important for *Shigella* treatment and disease control, and the World Health Organization (WHO) currently recommends ciprofloxacin (fluoroquinolone) as a first-line treatment, followed by pivmecillinam (beta-lactam), ceftriaxone (cephalosporin), and azithromycin (macrolide) as alternatives ^7^.

Fluoroquinolones are well-tolerated and were highly effective for shigellosis until resistance began to emerge in the early 2000s with sporadic cases of ciprofloxacin-resistant (cipR) *S. dysenteriae, S. flexneri*, and *S. boydii* being detected in India, Nepal, Pakistan, China, and Vietnam ^8–11^. Concurrently, *S. sonnei* with reduced susceptibility to fluoroquinolones became common across Asia ^6^, and fully cipR *S. sonnei* were characterized in India and Nepal soon after 9,12. These organisms carried the classical chromosomal point mutations in the quinolone resistance determining regions (QRDRs) at codon 83 (serine to leucine) and codon 87 (aspartic acid to glycine/asparagine) in *gyrA*, and at codon 80 (serine to isoleucine) in *parC* ^11,12^. From 2010 onwards, cipR *S. sonnei* emerged as a major global health concern, becoming widely distributed through international travel and ensuing domestic transmission ^13–15^. To date, cipR *S. sonnei* has been reported in children across Asia ^16,17^, as well as homeless individuals and men-who-have-sex-with-men (MSM) in Canada, the US, Taiwan, and the UK ^14,18–20^. Of further concern is the observation that cipR *S. sonnei* have the ability to acquire resistance to second-line alternative drugs such as ceftriaxone ^21^, which further limits alternative treatment options.

Our previous work demonstrated that all cipR *S. sonnei* globally were clonal and emerged once from a single lineage that likely arose in South Asia ^22^. Here, we describe the expansion of a single lineage III clade of cipR *S. sonnei* by providing phylo-temporal insights into its extant clonal dynamics using clinical samples obtained from children admitted to three paediatric facilities in Ho Chi Minh City (HCMC) between 2014 and 2016. We observe replacement of the resident ciprofloxacin susceptible (cipS) clone by the novel cipR lineage, as well as the concurrent and independent acquisition of a diverse range of resistance plasmids, which lead to MDR and XDR phenotypes. Through a detailed analysis of the plasmid content from commensal *E. coli* and *S. sonnei* isolated from a single patient and a series of conjugation experiments, we provide compelling evidence that ciprofloxacin exposure influences the *de novo* acquisition of MDR and XDR plasmids. Our data suggest that following the current international guidelines for *Shigella* therapy may lead to these cipR variants becoming resistant to alternative antimicrobials.

## Methods

### Study Design

The *S. sonnei* used in this study were isolated during a 2-year observational study conducted at three tertiary hospitals (Children’s Hospital 1, Children’s Hospital 2, and the Hospital for Tropical Diseases) in HCMC, Vietnam, between May 2014 and April 2016, as previously described (Supplementary Table 1) ^23^. In brief, children aged <16 years admitted to one of the three study hospitals with diarrhea (defined as ≥3 passages of loose stools within 24 hours) and >1 loose stool containing blood and/or mucus were recruited. A fecal sample was collected and processed within 24 hours after enrolment. All hospitalized patients received standard of care treatment at each of the study sites. Treatment and clinical outcomes (e.g. patient recovery status (three days after enrolment) and duration of hospital stay) were recorded by clinical staff at the study sites. For the phylogenetic analyses, we additionally included two cipR *S*. sonnei isolated from children attending the Hospital for Tropical Diseases in HCMC in October 2013.

Ethical approval was provided by the ethics committees of all three participating hospitals in HCMC and the University of Oxford Tropical Research Ethics Committee (OxTREC No.1045-13). Written consent from parents or legal guardians of all participants was obtained prior to enrolment.

### Microbiological methods

Fecal samples were inoculated onto MacConkey agar (MC agar; Oxoid) and xylose-lysine-deoxycholate agar (XLD agar, Oxoid) and incubated at 37°C. Non-lactose fermenting colonies grown on MC agar and/or XLD agar were sub-cultured on nutrient agar and identified biochemically (API20E, Biomerieux). Serological identification was performed by slide agglutination with somatic (O) antigen grouping sera following the manufacturer’s recommendations (Denka Seiken). Additionally, colony sweeps from MC agar were collected and suspended in 20% glycerol and stored at −80°C for further characterization.

Antimicrobial susceptibility testing was initially performed by the Kirby-Bauer disc diffusion method against ampicillin, chloramphenicol, trimethoprim-sulfamethoxazole, tetracycline, nalidixic acid, ciprofloxacin, azithromycin, gentamicin, amikacin, imipenem, and ceftriaxone. Subsequently, minimal inhibitory concentrations (MICs) against ciprofloxacin, azithromycin, gentamicin, and ceftriaxone were measured using the E-test (AB Biodisk), according to the manufacturer’s instructions. All antimicrobial testing was performed on Mueller-Hinton agar and susceptibility criteria were interpreted following the CLSI 2016 guidelines ^15^. MDR was defined as acquired non-susceptibility to at least one agent in three or more antimicrobial categories; XDR was defined as non-susceptibility to at least one agent in all but two or fewer antimicrobial categories (i.e. bacterial isolates remain susceptible to only one or two categories) ^24^. Detection of Extended Spectrum Beta Lactamase (ESBL) activity was performed for all isolates that were resistant to ceftriaxone using the combination disc method (cefotaxime, 30µg; ceftazidime, 30µg; with and without clavulanic acid, 10µg). ESBL-producing organisms were defined as those exhibiting a >5mm increase in the size of the zone of inhibition for the beta-lactamase inhibitor combinations in comparison to a third-generation cephalosporin without the beta-lactamase inhibitor.

### Isolation of commensal bacteria

For the purposes of this study we defined commensal organisms as organisms isolated from the stool samples of children thought not to be associated with the observed episode of diarrheal disease. In children with and without *Shigella* infections (i.e. symptomatic and asymptomatic children), non-*Shigella* organisms grown on MC plates were considered to be commensals. Colony sweeps from MC agar from *Shigella*-infected children were serially diluted and plated onto the MC agar without antimicrobial selection. All single colonies with a different color and morphology from that of *S. sonnei* were harvested, identified and homogenized in 20% glycerol. Subsequently, DNA was extracted from these commensal bacteria by boiling and was then subjected to qualitative real-time PCR with primers and probes specific for *Shigella* to detect if these samples were contaminated with *Shigella* ^25^. To determine the AMR gene and plasmid diversity in human commensal bacteria, we also included a subset of commensal bacteria recovered from the rectal swabs of 18 healthy children enrolled in a longitudinal cohort study with active surveillance for diarrheal disease in HCMC between 2014 and 2016 ^26^.

### Whole genome sequencing (WGS)

Genomic DNA from *S. sonnei* isolates and commensal bacteria was extracted using the Wizard Genomic DNA Extraction Kit (Promega, Wisconsin, USA) following the manufacturer’s recommendations. 50ng of genomic DNA from each sample was subjected to library construction using a Nextera kit, followed by whole genome sequencing on an Illumina MiSeq platform (Illumina, CA, USA) to generate 150 bp paired-end reads. Raw sequence data are available in the European Nucleotide Archive (project: PRJEB30967).

SNP calling for *S. sonnei* was performed as previously described ^6^. In brief, raw Illumina reads were mapped against *S. sonnei* reference genome Ss046 chromosome (accession number NC_007382) and virulence plasmid pSs046 (NC_007385.1) using SMALT version 0.7.4 (http://www.sanger.ac.uk/resources/software/smalt/). SNPs were called against the reference sequence and filtered using SAMtools ^27^. The allele at each locus in each isolate was determined by reference to the consensus base in that genome using SAMtools *mpileup* and removal of low confidence alleles with consensus base quality ≤20, read depth ≤5 or a heterozygous base call. SNPs occurring in non-conserved regions including prophages or repetitive sequences were removed. Subsequently, Gubbins ^28^ was used to identify recombinant regions from the whole genome alignment produced by SNP-calling isolates, and SNPs detected within these regions were also removed, resulting in a final set of 1,219 chromosomal SNPs.

### Phylogenetic analyses

The best-fit evolutionary model for the SNP alignment of all *S. sonnei* isolates was determined based on the Bayesian Information Criterion in jModelTest implemented in IQ-TREE software ^29^. A maximum likelihood phylogeny was subsequently reconstructed using IQ-TREE under the best-fit model (TVM). Support for the maximum likelihood tree was assessed via 1,000 pseudo-replicates. To explore the temporal signal in the data, the relationship between genetic divergence and date of sampling was estimated by using TempEst ^30^ to perform a linear regression analysis of root-to-tip distances, taken from the maximum likelihood tree, against the year of isolation (Supplemental Figure 1). Temporal phylogenetic inference was then performed using Bayesian Markov Chain Monte Carlo (MCMC) implemented in BEAST software (version 1.8.4) ^31^. For BEAST analysis, we first identified best-fit model combinations by performing multiple BEAST runs using the TVM nucleotide substitution model with constant, exponential growth or Bayesian skyline ^32^ demographic models, in combination with either a strict or a relaxed molecular clock (uncorrelated lognormal distribution) ^33^. For each BEAST run, path sampling and stepping-stone sampling approaches ^34,35^ were used to obtain the marginal likelihood estimates for model comparison. Bayes factor (the ratio of marginal likelihoods of two models) comparisons indicated that the TVM substitution model with a relaxed lognormal molecular clock and Bayesian skyline demographic model was the best fit to the data (Bayes factor >200). The standard deviation (SD) of inferred substitution rates across branches was 0.3 (95% highest posterior probability (HPD) = 0.14-0.47), providing additional support for a relaxed molecular clock. For the final analyses, we performed three independent runs with the best-fit model using a continuous 150 million generation MCMC chain with samples taken every 15,000 generations, and parameter convergence (indicated by effective sample size values >500) was assessed in Tracer (version 1.7). LogCombiner (version 1.8) was used to combine triplicate runs, with removal of 10□% burn-in.

### Resistome analysis of S. sonnei and commensal enterobacteria

From raw Illumina reads of *S. sonnei* and commensal enterobacteria Short Read Sequence Typer-SRST2 ^36^ was used to identify the acquired resistance genes and their precise alleles using the ARG-Annot database ^37^, as well as the plasmid replicons using the PlasmidFinder database ^38^. Multilocus sequence typing (MLST) of IncI plasmids ^39^ was also determined using SRST2. Raw Illumina reads were *de novo* assembled using Velvet, with the parameters optimized by Velvet Optimizer ^40^. Contigs <300 bp in size were discarded from further analyses and assembled contigs were annotated with Prokka ^41^. For the assembled sequences of *S. sonnei*, ABACAS ^42^ was used to map all the contigs against a concatenated reference sequence containing *S*. sonnei Ss046 chromosome (NC_007382), virulence plasmid pSs046 (NC_007385.1) and three small plasmids commonly found in *S. sonnei* belonging to global lineage III: spA (NC_009345.1), spB (NC_009346.1), spC (NC_009347.1). The unmapped assembled sequences presumably containing the *bla*CTX-M/*mphA* plasmid were subjected to manual investigation using BLASTN searching with the plasmid sequences available in GenBank, and comparative analysis was performed and visualized using ACT ^43^. For the assembled sequences of commensal bacteria carrying IncI and IncB/O plasmids, ABACAS was used to map contigs against full-length sequences of IncI and IncB/O plasmids identified in cipR *S. sonnei* and sequence comparisons were visualized using ACT. Nucleotide sequence homology between the mapped contigs and reference plasmid was subsequently identified using BLASTN. Taxonomic labels of each pooled sample of commensal bacteria were assigned using Kraken, a k-mer based classification tool ^44^.

### Plasmid profiling

Crude plasmid extractions from all *S. sonnei* isolates was performed using a modified Kado and Liu method ^45^. The resulting plasmid DNA was subjected to electrophoresis in 0.7 % agarose gel at 90 V for 3 hours, stained with ethidium bromide and photographed. *E. coli* strain 39R861 containing four plasmids with known sizes (7 kb, 36 kb, 63 kb, and 147 kb) was used as a marker. Plasmid profiles were compared using Bionumerics v5.1 software (Applied Maths, Austin, TX).

### ESBL plasmid digestion and sequencing

*E. coli* transconjugants resulting from conjugation between an ESBL-positive *S. sonnei* isolate and *E. coli* J53 (sodium azide resistance) were subjected to plasmid extraction using a plasmid Midi kit (Qiagen). For plasmid digestion, 500ng of each extracted plasmid DNA was digested with EcoRI enzyme (10 U/μl) (Fermentas), followed by electrophoresis on 0.8% agarose gel at 100V for 4 hours with 1 kb plus DNA ladder (Invitrogen). Plasmid restriction patterns were compared, and cluster analysis was performed using the UPGMA method and Jukes-Cantor correction using Bionumerics v5.1 software. For plasmid sequencing, 50ng of each plasmid DNA was subjected to library construction with a Nextera kit and sequenced using the MiSeq Illumina platform to generate 2×250 bp paired-end reads. *De novo* assembly was subsequently performed using SPADES v3.11 ^46^ and assembled contigs were annotated using Prokka v1.11 ^41^.

### Nanopore sequencing

Plasmid DNA extracted from the commensal *E. coli* carrying the IncI/blaCTX-M-15 plasmid was initially sequenced using Illumina MiSeq to generate 2×250bp paired-end reads (accession number: ERS3050916). However, the *de novo* assembly failed to produce a complete plasmid sequence. To improve the plasmid assembly, we then performed a single run on a MinION to generate longer reads. For MinION library preparation and sequencing, we used the rapid 1D sequencing kit SQK-RAD001 (Oxford Nanopore Technologies, Oxford, UK), following the manufacturer’s recommendations. We used the MinION Mk1 sequencer, FLO-MIN106 flow cell and MinKNOW software v1.1.20 for sequencing, and protocol script NC_48Hr_Sequencing_Run_FLO-MIN106_SQK-RAD001_plus_Basecaller.py for local base-calling. MinION reads were converted from fast5 to fastq format using the script fast52fastq.py. SPADES version 3.11 was subsequently used to produce a hybrid assembly of MinION data and Illumina data. Raw MinION reads were deposited in ENA (accession number: ERS3050922).

### Bacterial conjugation

Bacterial conjugation was first performed between each of the 40 *bla*_CTX-M_/*mphA*-carrying *S. sonnei* isolates associated with all the plasmid acquisitions and *E. coli* J53 (sodium azide resistant) by combining equal volumes (5mL) of overnight Luria-Bertani (LB) cultures. Bacteria were conjugated for 12 hours in LB broth at 37°C and *E. coli* transconjugants were selected on medium containing sodium azide (100 mg/l) plus ceftriaxone (6 mg/l) or sodium azide (100 mg/l) plus azithromycin (24 mg/l). To measure plasmid transfer from commensal *E. coli* to cipR *S*. sonnei and investigate the effect of ciprofloxacin on the conjugation efficiency, we first screened commensal *E. coli* isolates for ESBL activity and ciprofloxacin susceptibility from the pooled colony sweeps on MC agar. Subsequently, bacterial conjugation was performed between each of the 13 cipS ESBL-positive commensal *E. coli* isolates (donor) and the cipR ESBL-negative *S. sonnei* 03-0520 (recipient) in LB broth with and without supplementation of ciprofloxacin (0.25, 0.5, 0.75 × MIC of the donor organism). Successful transconjugants were selected on MC agar containing ciprofloxacin (4 mg/l) and ceftriaxone (6 mg/l). For all conjugation experiments, the conjugation frequency was calculated as the number of transconjugants per recipient cell.

## Results

### The development of an XDR phenotype in ciprofloxacin-resistant Shigella sonnei

Between January 2014 and July 2016, we isolated 79 *S. sonnei* from children hospitalized with dysenteric diarrhea in our study sites; 75.9% (60/79) of these were cipR. A time-scaled phylogenetic reconstruction demonstrated that all except one cipR *S. sonnei* comprised a distinct clade, which was distantly related to the ciprofloxacin-susceptible (cipS) isolates (Figure 1). The most recent common ancestor (MRCA) of the cipR clade in Vietnam was estimated to date back to late 2008 (95% HPD; 2007.1 – 2010.3) – several years prior to the first known cases of cipR *S. sonnei* in Vietnam, which was detected in HCMC in October 2013. The phylogeny depicted a clonal expansion from a single cipR organism that we have previously shown to have originated in South Asia and disseminated internationally ^22^. All organisms within the cipR clade were classical triple mutants (*gyrA*-S83L, *gyrA*-D87G, and *parC*-S80I) conferring high-level ciprofloxacin resistance (MIC ≥8 μg/ml). Conversely, isolates belonging to the resident cipS *S. sonnei* clade harbored only a single mutation in *gyrA* (either S83L (16 isolates) or D87Y (4 isolates)).

**Figure 1.**
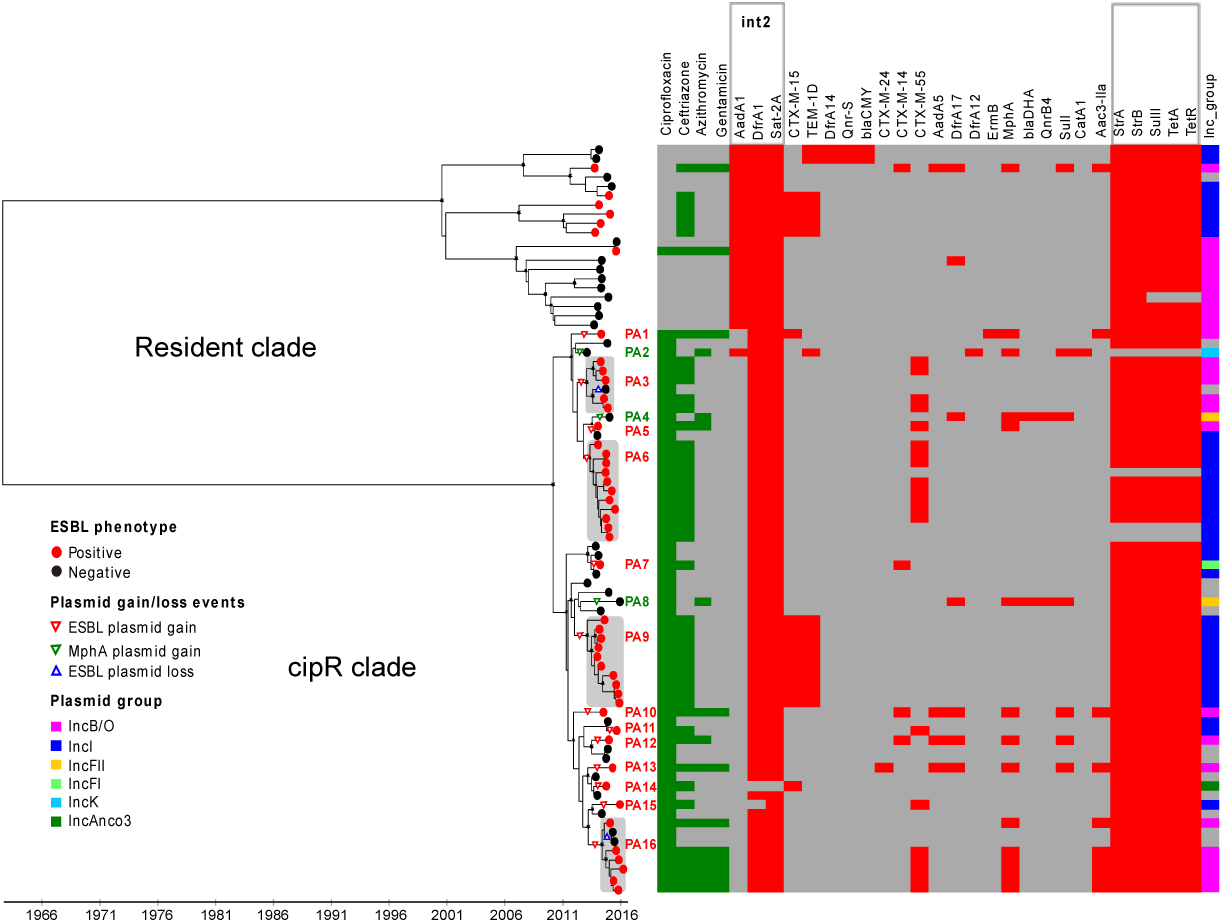
The temporal phylogenetic structure of *Shigella sonnei* in Vietnam, 2014 and 2016 Maximum clade credibility phylogeny showing two distinct clades corresponding to the resident and the cipR *S. sonnei* populations. The black asterisks indicate posterior probability support ≥70% on internal nodes. The red circles at terminal leaves highlight the ESBL-positive isolates. Triangles indicate plasmid gain/loss events. Sixteen plasmid acquisitions (PAs) were reconstructed across the tree and designated as PA1-PA16. The columns on the right correspond to: the resistance phenotype to key antimicrobials (green), the presence of AMR genes (red), and the presence of different plasmid groups (multiple colors), respectively.

Using BEAST, we estimated the median substitution rate of the *S. sonnei* population to be 8.2×10^−7^ substitutions base ^−1^ year ^−1^ (95% highest posterior density (HPD); 5.9×10^−7^ to 10.8×10^−7^), which is comparable to previous estimates of the mutation rate within the resident cipS *S. sonnei* population in Vietnam ^47^. Additionally, the cipR isolates exhibited a substantially lower median pairwise SNP distance (15 SNPs, IQR: 10-20 SNPs) than the resident cipS *S. sonnei* isolates (96 SNPs, IQR: 69-111 SNPs), providing strong evidence of a more recent importation or expansion. These data suggest that cipR *S. sonnei* underwent a rapid clonal expansion and successfully persisted, displacing the resident cipS *S. sonnei* as the dominant *S. sonnei* lineage circulating in the human population of southern Vietnam.

During the sampling period, the proportion of cipR *S. sonnei* increased significantly from 60.7% (17/28) in 2014 to 93.8% (15/16) in 2016 (*p=*0.01; Chi-squared test). Almost all isolates carried AMR genes on a chromosomally integrated class II integron (*dfrA1, sat-2A*) and a small spA plasmid (*strAB, sulII, tetAR*) encoding resistance to tetracycline, streptomycin, and co-trimoxazole. Notably, during their circulation in southern Vietnam, the cipR *S. sonnei* acquired resistance to further important antimicrobial classes, including third-generation cephalosporins, macrolides, and aminoglycosides, consequently creating XDR variants. In 2014, the proportion of co-resistance against ceftriaxone, ceftriaxone-azithromycin, and ceftriaxone-azithromycin-gentamicin was 59% (10/17), 11.8% (2/17), and 5.9% (1/17), respectively. These respective proportions increased to 87% (13/15), 47% (7/15), and 40% (6/15) in 2016.

Our analyses show that co-resistance in cipR *S. sonnei* was generated by sustained and independent acquisitions of ESBL-encoding plasmids (plasmid acquisition events, herein referred to as PAs, Figure 1). These plasmids carried differing variants of the *bla*_CTX-M_ gene and/or *mphA*. Notably, this phenomenon was not characterized by selection of the same plasmid/clone combination; we identified at least thirteen independent acquisitions of ESBL-encoding plasmids across the phylogenetic tree (Figure 1). Plasmids of incompatibility groups IncB/O and IncI were the most common vehicles associated with *bla*_CTX-M_.

More specifically, we found that IncB/O plasmids were independently acquired on at least seven occasions; these plasmids carried an array of ESBL genes, including *bla*_CTX-M-55_ (PA3, 5, 16), *bla*_CTX-M-14_ (PA10, 12), *bla*_CTX-M-15_ (PA1), and *bla*_CTX-M-24_ (PA13). Critically, in 6/7 IncB/O PAs, the *bla*_CTX-M_ gene was associated with *mphA* and *acc6-IIa* genes, leading to an XDR phenotype additionally encompassing resistance to third-generation cephalosporins, macrolides, and aminoglycosides. Similarly, IncI plasmids were acquired on four independent occasions and carried only a *bla*_CTX-M_ gene (*bla*_CTX-M-15_ (PA9) and *bla*_CTX-M-55_ (PA6, 11, 15)). Furthermore, *bla*_CTX-M_ genes were acquired on IncFI (*bla*_CTX-M14_ (PA7)) and IncAnco3 (*bla*_CTX-M-15_ (PA14)) plasmid backbones. Three cipR non-ESBL isolates also acquired an *mphA* gene associated with IncFII (PA4, 8) and IncK (PA2) plasmids.

The acquisition of a resistance plasmid was sporadically followed by continued circulation and geographical expansion of the resistant clone, as observed for the IncB/O/*bla*_CTX-M-55_ (PA3), IncB/O/*bla*_CTX-M-55_*-mphA-aac6-IIa* (PA16), IncI/*bla*_CTX-M-55_ (PA6), and IncI/*bla*_CTX-M-15_ plasmids (PA9) (Figure 1). The inferred time from the most recent common ancestor to the youngest isolate in each resistant clone was estimated to be at least three years. We also detected the loss of IncB/O plasmids on two occasions, suggesting a potential lack of IncB/O plasmid stability in comparison to the IncI plasmids.

### The structure of XDR plasmids in ciprofloxacin-resistant Shigella sonnei

We assessed the plasmid content of all *S. sonnei* isolates by comparing the banding patterns of crude undigested plasmid extracts. Our data showed that all cipR isolates exhibited a distinct plasmid profile from that of the resident Vietnamese *S. sonnei* isolates (Supplementary Figure 2). Additionally, all cipR *S. sonnei* isolates carrying *bla*_CTX-M_ and/or *mphA* consistently harbored a large (90 kb to 110 kb) plasmid. An analysis of the EcoRI digestion profiles of these ESBL-encoding plasmids showed two major independent clusters, consistent with IncI and IncB/O plasmid backbones (Figure 2A). Notably, the genetic structure within each plasmid group appeared to be highly conserved, with the IncI and IncB/O plasmids sharing ∼70% and ∼60% similarity in their respective restriction patterns.

**Figure 2.**
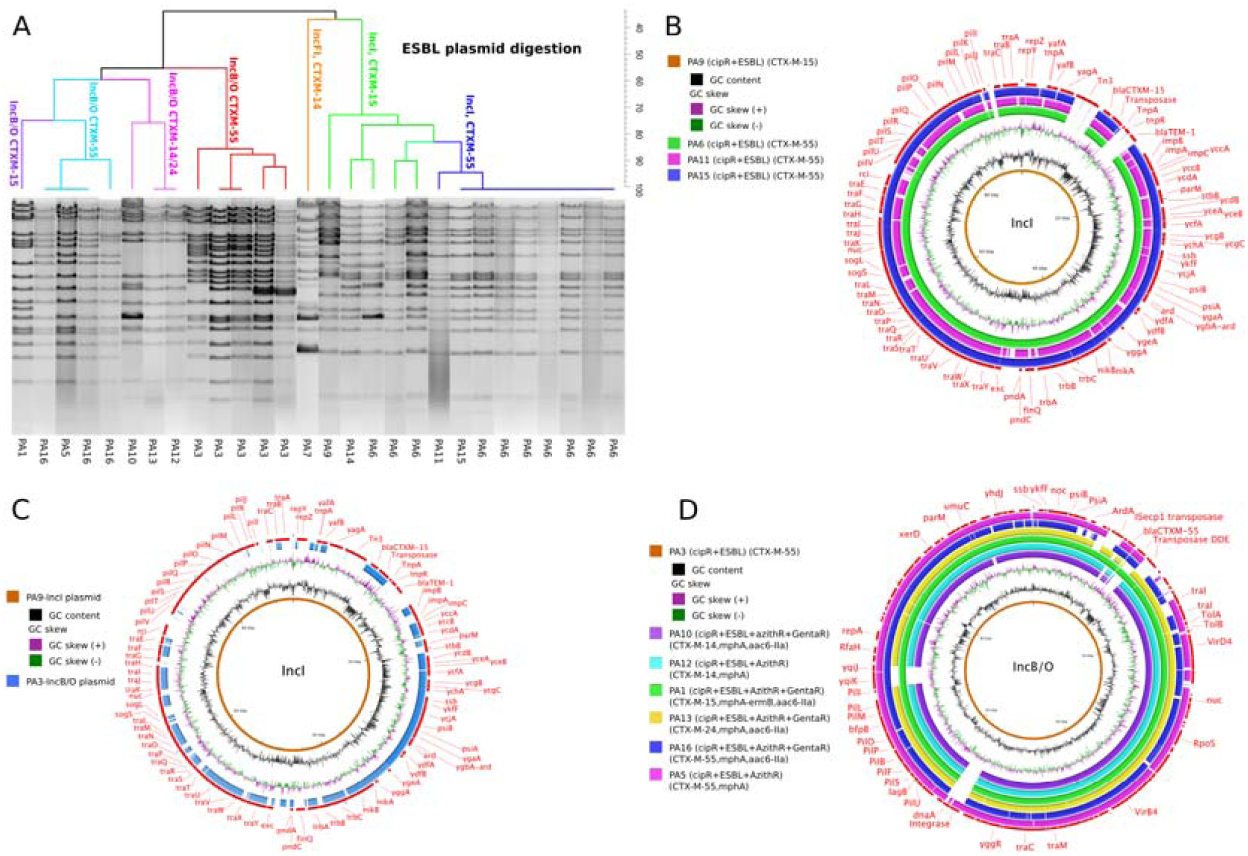
The Diversity of ESBL-encoding plasmids in ciprofloxacin-resistant *Shigella sonnei* A) Dendrogram showing the similarities in restriction digestion patterns of ESBL-encoding plasmids associated with independent plasmid acquisitions. B) BLAST comparisons of IncI plasmids associated with four independent acquisitions (PA6, 9, 11, 15). The central circle is the full reference sequence of the IncI plasmid associated with PA9, with similarity between the reference sequence and other IncI plasmids shown as concentric rings. C) BLASTN comparison between IncI and IncB/O plasmid structures, in which the central circle is the IncI plasmid (PA9). D) BLAST comparisons of IncB/O plasmids associated with seven independent acquisitions (PA1, 3, 5, 10, 12, 13, 16). The central circle is the full reference sequence of the IncB/O plasmid associated with PA3, with similarity between the reference sequence and other IncB/O plasmids shown as concentric rings.

Additional plasmid sequencing and comparative analyses found that the IncI plasmids, acquired on four occasions (PA6, 9, 11, 15), shared a conserved backbone of ∼84 kb (coverage: 80-100%, nucleotide identity: 99-100%). This conserved region contained typical structures associated with self-transmissible IncI plasmids, including a type IV *pil* operon (*pilI-PilV*), *traABC* regulatory genes, the *tra/trb* type IV secretion system genes, the origin of transfer (*oriT* including *nikA* and *nikB*), and conjugal leading region (*ssb, psiA-psiB, parB* homolog, *ardA*). The IncI/*bla*_CTX-M-15_ plasmid belonged to sequence type 16 (ST16) and was nearly identical to the previously described *S. sonnei* IncI plasmid pKHSB1 (accession number: NC_020991), which has been maintained in the resident Vietnamese *S. sonnei* population since 2006 ^47^. Alternatively, the IncI/*bla*_CTX-M-55_ plasmids belonged to ST167 (one allele different from ST16) and did not harbor the Tn3 transposon-mediated IS*ecp1*-*bl*aCTX-M-15, but had an insertion of IS*ecp1*-*bla*CTX-M-55 between *yagA* and *yafB* (Figure 2B).

The IncB/O plasmids, acquired on seven occasions (PA1, 3, 5, 10, 12, 13, 16), also shared a conserved genetic structure of ∼90 kb (coverage: 75-100%, nucleotide identity 99-100%) (Figure 2D). In comparison to plasmid sequences in GenBank, our IncB/O plasmid backbone shared the highest similarity with IncB/O plasmids from an *E. coli* (pECAZ161, accession number: CP19011), a *S. flexneri* (pSF150802, accession number: CP030917.1) and an *S. sonnei* (p866, accession number: CO022673.1); the overall synteny ranged from 72% to 94%, with 99% sequence identity. The IncB/O plasmid backbone contained comparable conjugative IncI plasmid modules; however, the *pil* operon (∼12 kb) exhibited extremely low sequence identity to that of the IncI plasmid (coverage 1%, identity 78%) (Figure 2C). The size of IncB/O plasmids varied from 95 kb to 110kb, depending on the complement of resistance gene cassettes. These plasmids carried a wide range of *bla*_CTX-M_ mobile elements, including IS*ecp1*-IS*5*-*bla*_CTX-M-55_, IS*66*-*bla*_CTX-_ M-55-*orf-tnpA*, IS*ecp1*-*bla*_CTX-M-15_-*orf-tnpA*, IS*5*-*bla*_CTX-M-14_-IS*ecp1*, and IS*ecp1b*-*bla*_CTX-M-24_. Additionally, these IncB/O plasmids also contained other transposable elements associated with *mphA* (IS*6-mphA-mrx-mphr*) and *aac6-IIa* (IS*4*-*aac6-IIa-tmrB*) adjacent to *bla*_CTX-M_-carrying elements. One IncB/O plasmid additionally carried the *ermB* gene associated with the ISCR3 family (ISCR3-*groEL-ermB-ermC*).

Aside from the two main IncI and IncB/O plasmid groups, one cipR *S. sonnei* isolate had gained a *bla*_CTX-M-14_-carrying IncFI plasmid, which was identical to a previously described plasmid (pEG356) from a Vietnamese *S. sonnei* isolate (accession number: FN594520); a further isolate acquired a phage-like IncAnco3 plasmid carrying *bla*_CTX-M-15_; three other isolates gained IncK and IncFII plasmids carrying the *mphA* gene cassette. The IncAnco3, IncK, and IncFII plasmids were most similar to described *E. coli* plasmids in GenBank, including pAnco1 (accession number: KY515224.1, coverage 91%, identity 98%), pEC1107 (accession number: MG601057.1, coverage 78%, identity 94%), and pEC105 (accession number: AY458016.1, coverage 59%, identity 100%), respectively.

### Commensal E. coli as a source of ESBL-encoding plasmids for ciprofloxacin-resistant Shigella sonnei in the human gut

Given the diversity of the AMR plasmids observed in cipR *S. sonnei*, their similarity to *E. coli* plasmids, and the fact that humans are the only natural reservoir for *S. sonnei*, we speculated that these plasmids had been transferred from commensal *E. coli* into *S. sonnei* during infection. Consequently, we performed additional characterization of AMR genes and plasmid diversity in commensal Enterobacteriaceae isolated from the same fecal samples that contained *S. sonnei* and from rectal swabs taken from healthy children. Metagenomic sequencing of these mixed bacterial populations (lacking *Shigella*) from MC plates indicated that *E. coli* was the most commonly isolated commensal Enterobacteriaceae (47/48 pooled colonies), followed by *Klebsiella pneumoniae* (7/48 pooled colonies) and *Enterobacter cloacae* (1/48 pooled colonies).

The resulting sequence data identified a substantial quantity of AMR genes and plasmid backbones in the commensal Enterobacteriaceae (Supplementary Figure 3 and Figure 3A). We observed a particularly high prevalence of CipR commensal Enterobacteriaceae; this has been observed previously and is considered to be associated with sustained antimicrobial exposure and competition in the gastrointestinal tract ^48^. Furthermore, a number of different AMR determinants were found to be present in both the commensal bacteria and the cipR *S. sonnei*. For example, *bla*_CTX-M_, *mphA, aac3-IIa*, and *ermB* were found to be present in cipR *S. sonnei* and 92% (44/48), 75% (36/48), 52% (25/48), and 38% (18/48) of pooled commensal Enterobacteriaceae, respectively (Figure 3A). IncF (IncFII, IncFIA, IncFIB, and IncFIC) plasmids were found to be the most prevalent replicon types in the commensal Enterobacteriaceae. However, we additionally identified IncI and IncB/O plasmids in commensal *E. coli* from the fecal samples of three children infected with *S. sonnei* and three healthy children, respectively.

**Figure 3.**
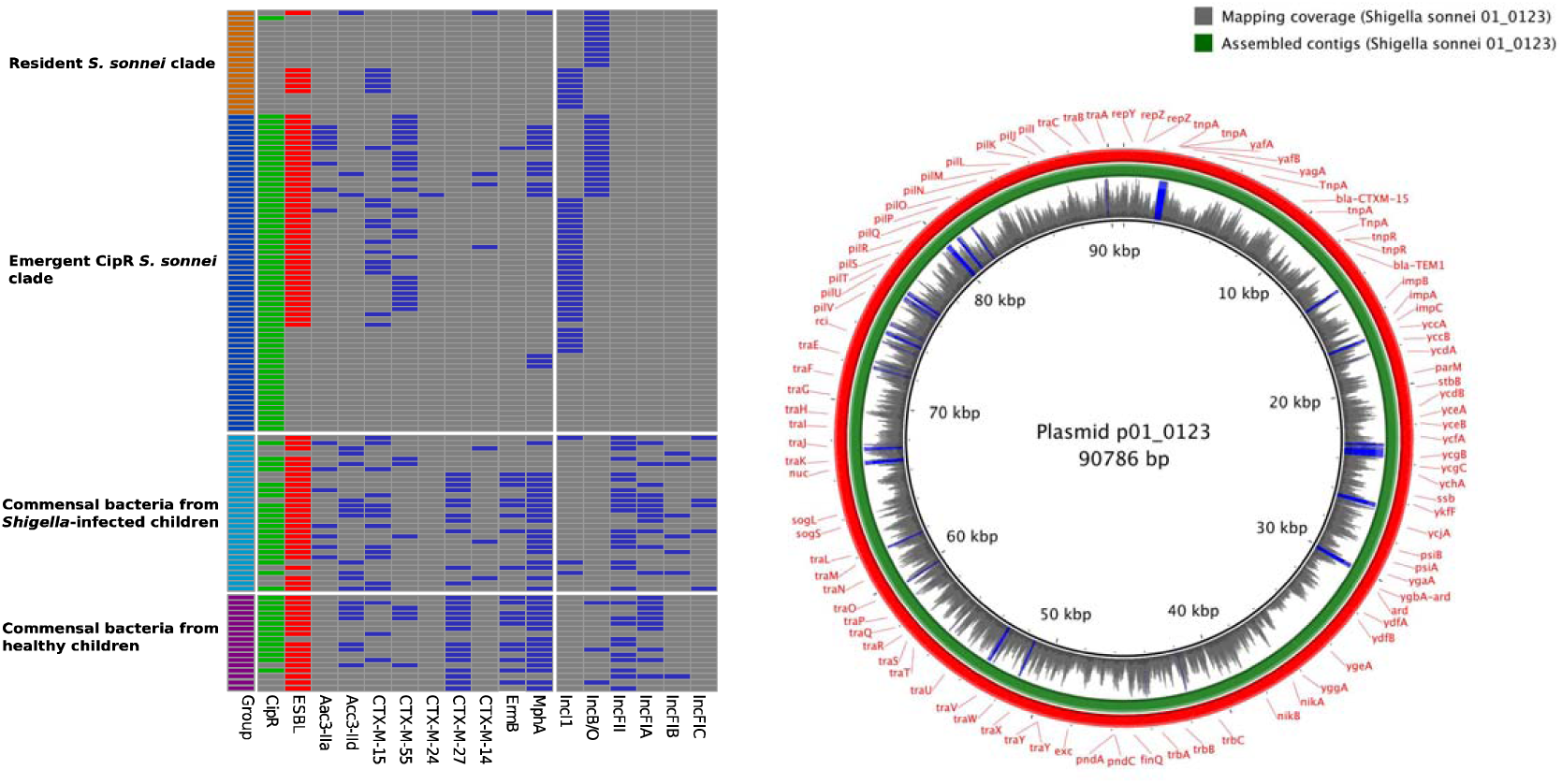
Antimicrobial resistance genes and plasmids in human commensal bacteria and *Shigella sonnei* A) The first column highlights the four different sample types. Fecal/rectal swab cultures with ciprofloxacin-resistant and ESBL-producing isolates are highlighted in green and red (second and third columns, respectively). The remaining columns show the presence of key antimicrobial resistance genes and plasmid groups (blue) in commensal bacteria and *S. sonnei*. B) The central circle is the full sequence of plasmid p01-0123 assembled from Nanopore sequences of commensal *E. coli*. The next ring shows the depth of coverage from raw Illumina reads of ciprofloxacin-resistant *Shigella sonnei* strain 01-0123 mapped onto the central reference sequence. Graph height is proportional to the number of reads mapping at each nucleotide position in the reference genome from 0 to 30x coverage. Regions with plasmid coverage greater than 30x are shown as solid blue bands. The green ring shows the BLASTN comparison between assembled sequences of *Shigella sonnei* strain 01-0123 and the central reference sequence. The red ring indicates the gene annotations of the central reference sequence.

We next aimed to identify comparable plasmid structures between *E. coli* and *S. sonnei*. The IncB/O plasmids found in commensal *E. coli* from three healthy children (subjects 22889, 22959, and 22274) exhibited high levels of sequence similarity to the IncB/O plasmid backbone acquired by cipR *S. sonnei* (coverage/identity: 76/99%, 94/99%, and 99/97%; respectively). Similarly, among the three commensal *E. coli* samples carrying IncI plasmids, we identified an IncI plasmid from a commensal *E. coli* without a Tn3 transposon-mediated *bla*_CTX-M-15_, which displayed high sequence similarity (coverage 81%, identity 98%) to an IncI plasmid from cipR *S. sonnei*. More significantly, a commensal *E. coli* originating from a patient infected with a cipR *S. sonnei* (01-0123) carried an analogous IncI/*bla*_CTX-M-15_ plasmid. We isolated a single ESBL-producing commensal *E. coli* from this MC plate and subjected the plasmid to long read Nanopore sequencing. The sequencing resulted in a 90,786 bp circularized plasmid sequence, harboring *bla*_CTX-M-15_ and *bla*_TEM1_ on a Tn3 transposon. The raw IncI plasmid sequence from the corresponding cipR *S. sonnei* 01-0123 was mapped against the commensal *E. coli* plasmid sequence and produced a plasmid with 100% coverage (mean mapping coverage: 15, standard deviation: 7). The assembled plasmid contigs from cipR *S. sonnei* 01-0123 shared 100% sequence identity (Figure 3B). These data and the location of this organism on the phylogenetic tree suggest that this resistance plasmid was potentially transferred *in vivo* between commensal *E. coli* and cipR *S. sonnei* 01-0123 in the gut of the child.

### Ciprofloxacin increases the conjugation frequency of ESBL plasmids between commensal Escherichia coli and ciprofloxacin-resistant Shigella sonnei

Our data illustrates that commensal *E. coli* are an important reservoir of AMR genes and may be transferred to *S. sonnei in vivo*. Furthermore, the high diversity of AMR plasmids observed here in a single *S. sonnei* lineage is atypical and has not been previously observed in a geographically restricted clonal expansion. The reason for this observation is unclear but we suspect is associated with the combination of a permissive circulating clone, exposure to fluoroquinolones, and a wide variety of AMR plasmids in the resident commensal population. We also observed that the majority of *S. sonnei* infected children (85%, 67/79) were treated with ciprofloxacin, an antimicrobial agent that can trigger the SOS response and promote horizontal gene transfer in bacteria ^49–54^. Consequently, we hypothesized that this array of resistance plasmids was associated with a cipR phenotype and that ciprofloxacin treatment may facilitate plasmid transfer *in vivo*.

To test this hypothesis, we first identified cipS/ESBL+ commensal *E. coli* donors and attempted to mobilize these plasmids into a cipR/ESBL-*S. sonnei* recipient. Screening identified that the majority of commensal *E. coli* (35/48) recovered from the MC plate sweeps were both cipR/ESBL+. The remaining 13 commensal *E. coli* isolates were cipS (MIC ≤ 1 mg/L)/ESBL+; nine were derived from children infected with *S. sonnei* and four from healthy children. ESBL plasmids from 9/13 of the commensal *E. coli* could be conjugated into the cipR/ESBL-*S. sonnei*. The conjugation frequencies were high, ranging from 4×10^−7^ to 1.6×10^−3^/recipient cells. The supplementation of 0.25× MIC ciprofloxacin into the conjugation media did not have a significant effect on the frequency of plasmid transfer. However, when the conjugation media was supplemented with 0.5x MIC ciprofloxacin (of the cipS *E. coli* donor), 4/9 commensal *E. coli* (22784, 01-0123, 02-1936, and 22959) demonstrated respective increases in conjugation frequencies of ESBL plasmids to cipR *S. sonnei* of 3, 6, 11, and 36-fold, in comparison to media without ciprofloxacin (Figure 4). These respective conjugation frequencies increased to 4, 10, 25 and 42-fold when the concentration of ciprofloxacin was increased to 0.75x MIC. Additionally, a single commensal *E. coli* isolate (22978) exhibited a 7-fold increase in conjugation frequency in medium supplemented with 0.75x MIC ciprofloxacin, despite this effect not being observed in media containing 0.5x MIC ciprofloxacin.

**Figure 4.**
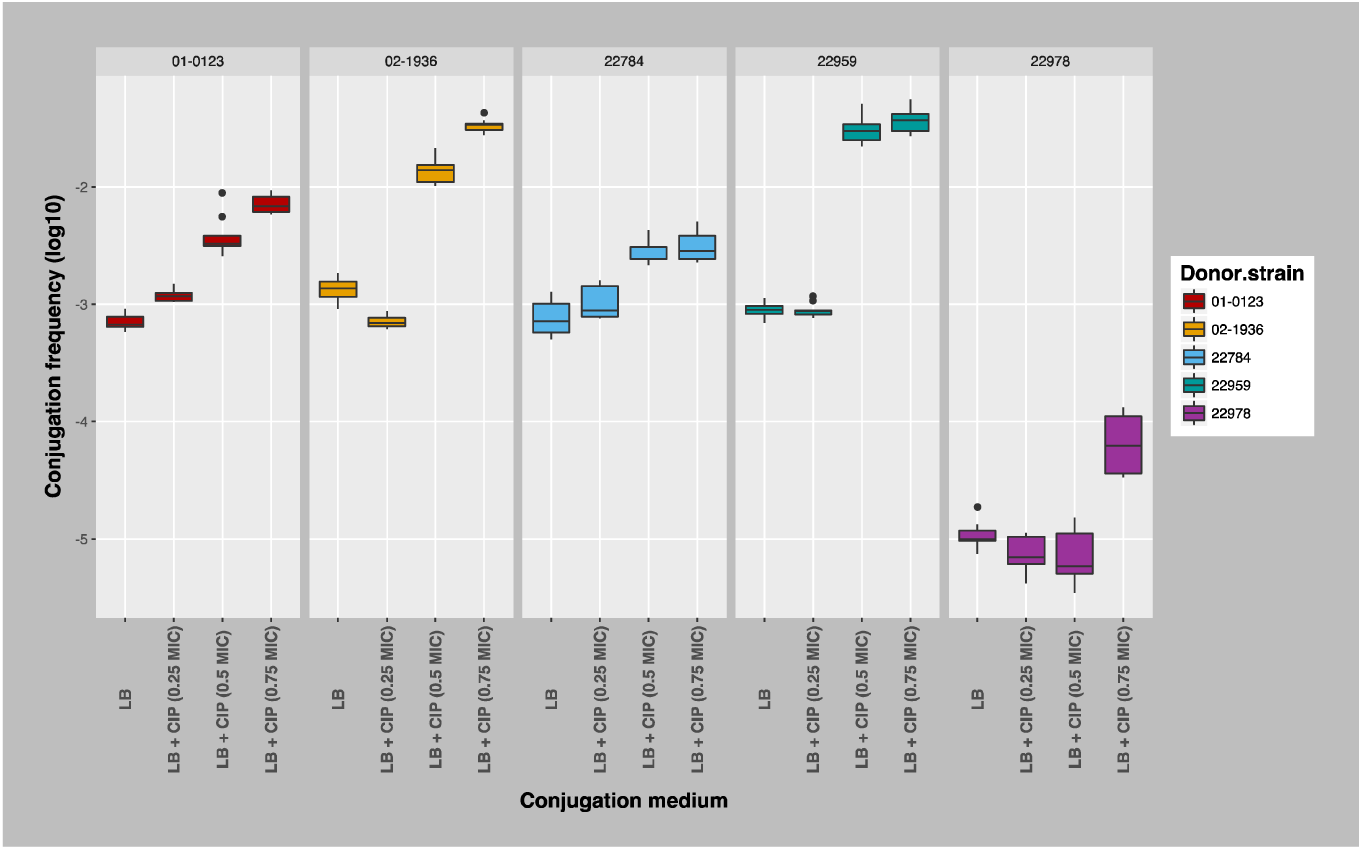
The Conjugation efficiency of ESBL-encoding plasmids from human commensal *E. coli* to ciprofloxacin-resistant *Shigella sonnei* with and without supplementation with ciprofloxacin Boxplots showing the conjugation frequencies (log10) between five commensal *E. coli* isolates and cipR *S. sonnei* strain 03-0520 with and without supplementing conjugation media with 0.25x, 0.5x, and 0.75x MIC of the donor cipS commensal *E. coli*. Each conjugation experiment was performed in triplicate in each condition.

Plasmid sequencing demonstrated that *E. coli* 01-0123 (ciprofloxacin MIC: 0.25 mg/L) carried an IncI/*bla*_CTX-M-15_ plasmid (as described above). *E. coli* 02-1936 (ciprofloxacin MIC: 0.016 mg/L) and 22784 (ciprofloxacin MIC: 0.38 mg/L) carried IncF/*bla*_CTX-M-27_ plasmids that shared high similarity to the IncF plasmid pC15 in Genbank (accession number: AY458016, ∼ 92 kb) (coverage: 75% and 85%, identity: 99% and 98%, respectively). *E. coli* 22959 (ciprofloxacin MIC: 0.38 mg/L) harbored an IncB/O/ *bla*_CTX-M-27_ plasmid exhibiting high genetic similarity to the IncB/O plasmid acquired by cipR *S. sonnei* as described above (coverage 94%, identity 99%). *E. coli* 22978 (ciprofloxacin MIC: 0.5 mg/L) carried an IncF/*bla*_CTX-M-27_ similar to the IncF plasmid pDA33135 in Genbank (accession number: CP029577.1, ∼ 139 kb) (coverage 94%, identity 99%).

## Discussion

Since its introduction in the 1980s, ciprofloxacin has become one of the most commonly used antimicrobials worldwide due to its low cost and clinical effectiveness against a wide range of Gram-positive and Gram-negative bacterial infections. The extensive use of ciprofloxacin in humans and animals inevitably led to a rapid increase in reduced susceptibility to ciprofloxacin in both Gram-negative and Gram-positive bacteria during the 1990s ^55^. Since the turn of the century, multiple cipR clones in various pathogenic species/serotypes have emerged and spread successfully in various countries, with many eventually disseminating internationally. Organisms with internationally successful cipR clones include *Salmonella* Typhi ^56^ and *Shigella dysenteriae* type 1 ^57^ in South Asia, and methicillin-resistant *Staphylococcus aureus* ST22 ^58^, ST131-H30 clone of *E. coli* ^58^, *Salmonella* Kentucky ST198 ^60^, *Clostridium difficile* 027 ^61^, *Shigella sonnei* ^22^ and *E. coli* ST1193 ^62–65^. Tracking the global transmission and local establishment of these clinically important clones through routine surveillance, particularly with the integration of genomics, has become essential for guiding public health control strategy and clinical practice. Here, by decoding the genomic sequences of *S. sonnei* isolated in Vietnam between 2014 and 2016, we provide unparalleled insight into the local clonal establishment and AMR dynamics of cipR *S. sonnei* as it entered a new human population. Our work outlines the progression and co-circulation of multiple XDR *S. sonnei* clones in Vietnam, some of which have gained resistance to all antimicrobial therapies currently recommended by WHO for the treatment of *Shigella*.

Although the cipR *S. sonnei* sublineage Central Asia III has spread internationally, the development of XDR within this lineage has not been reported previously. *S. sonnei* are highly efficient at spreading internationally; therefore, the identification and pervasiveness of AMR in these organisms means that future investigations should monitor their international circulation to provide early warning for public health authorities and healthcare providers. More specifically, the emergence and expansion of cipR XDR *S. sonnei* clones associated with IS*ecp1*-*bla*_CTX-M-55_ raises a major concern regarding the epidemic potential of this novel CTX-M ESBL variant in *S. sonnei* ^66–68^. A significant burden of shigellosis, high prevalence of AMR among Gram-negative commensal bacteria, and the purchasing of antimicrobials in the community may be factors contributing to the emergence and maintenance of XDR *S. sonnei* in Vietnam.

We describe multiple different plasmid structures in the cipR *S. sonnei* population, distinguishing their dynamics from those of the resident *S. sonnei* population, which underwent a clonal expansion characterized by a single plasmid structure ^47^. In most cases when a *mphA*/*bla*_CTX-M_ plasmid was acquired, the plasmids appeared not to become established in the population. This observation suggests that these structures could have a sustained fitness disadvantage in the absence of antimicrobial pressure. Conversely, the successful maintenance of four independent XDR *S. sonnei* clones warrants further investigation into their potential fitness, plasmid stability, and future evolutionary trajectories. Additionally, the selected IncB/O and IncI ESBL-encoding conjugative plasmids acquired and maintained by cipR *S. sonnei* suggest plasmid preferences in this species. IncI and IncB/O plasmids belong to the IncI-complex (IncI, IncB/O, IncK, IncZ), which have comparable antisense RNA plasmid replication control mechanisms ^69^. Our results concur with previous reports that proposed a commonality of IncI-complex plasmids associated with the *bla*_CTX-M_ element in *S. sonnei* from countries at various stages of economic development ^21,47,70–74^. The reasons for these specific plasmid-host combinations remain elusive; however, we found that IncI and IncB/O plasmids displayed the highest *in vitro* conjugation efficiencies compared to other ESBL-encoding plasmids. Moreover, the IncB/O plasmid group was found to be the second most commonly identified plasmid group in commensal and pathogenic *E. coli* from humans and animals ^75^, while the conjugative IncI group was can also be highly prevalent in commensal bacteria from infants ^76^. The regular sampling of these plasmids by cipR *S. sonnei* could be attributed to several factors, including the close genetic relatedness between *Shigella* and *E. coli*, the propensity of *S. sonnei* to acquire AMR plasmids, and the circulation of highly transmissible AMR plasmids in commensal *E. coli*. To combat the emergence and circulation of AMR Gram-negative bacteria, a better understanding of plasmid-host interactions, plasmid stability, and the role of plasmids in the fitness of cipR *S. sonnei* are now critical.

The routine acquisition of a wide variety of ESBL-encoding plasmids by cipR *S. sonnei* reflects extensive interspecies gene flow from a substantial local gene pool, possibly as a result of bacterial response to selective pressures exerted by widespread and largely uncontrolled antimicrobial use. These plasmids appear to originate from bacterial hosts that share the same ecological niche as *S. sonnei*. We performed WGS of selected commensal bacteria from fecal samples infected with *S. sonnei* and from healthy children and identified an extensive range of AMR genes/plasmids that conferred resistance to all antimicrobial classes in these commensal organisms. This diversity included the IncI and IncB/O plasmids that had been routinely acquired by cipR *S. sonnei*. We also provide evidence for *in vivo* IncI/*bla*_CTX-M-15_ plasmid transfer between commensal *E. coli* and cipR *S. sonnei* in a single patient; however, we cannot resolve the directionality of plasmid movement or discount the role of other components of the human microbiome as the original donor of this plasmid. Potential plasmid transfer between commensal Gram-negative bacteria and *Shigella* spp. in the human gut has been suggested previously ^77,78^. The large biomass of Enterobacteriaceae in the human gastrointestinal tract and the apparent common circulation of IncI and IncB/O plasmids in commensal bacteria in the gastrointestinal tracts of Vietnamese children suggests that the direction of plasmid transfer is more likely to be from commensal bacteria (potentially *E. coli*) to *S. sonnei*.

We additionally aimed to assess the role of ciprofloxacin in facilitating the transfer of ESBL-encoding plasmids from commensal bacteria to cipR *S. sonnei*. Our data show that exposure to sub-[inhibitory concentrations to ciprofloxacin may facilitate the transfer of ESBL-encoding plasmids between commensal *E. coli* and cipR *S. sonnei*. As the majority of commensal *E. coli* were cipR, our results suggest that horizontal plasmid transfer between cipR Gram-negative organisms and *Shigella* may occur at higher frequencies in the presence of increasing ciprofloxacin concentrations. Our observations question the effect of ciprofloxacin on the composition of commensal bacterial and transfer dynamics of AMR determinants in the human gut during and after treatment. In the clinical study in which the *S. sonnei* described here were isolated, the majority of *S. sonnei* infected children (85%, 67/79) were treated empirically with ciprofloxacin. We found that the clinical outcomes (duration of hospitalization) between children infected with cipR (51 cases) versus cipS *S. sonnei* (16 cases) were comparable (median 4 days (IQR: 3-6.5 days) versus 3 days (IQR: 2-4)) ^79^. Additionally, four cases (three infected with cipR *S. sonnei*) did not receive antimicrobial treatment but still recovered in a similar time period. These supporting data call for a re-evaluation of the necessity and benefit of treating children with *S. sonnei* dysentery with ciprofloxacin. Treatment with fluoroquinolones in the absence of appropriate diagnostics and susceptibility testing could potentially select for the maintenance and transmission of XDR *S. sonnei* and promote horizontal plasmid transfer between commensal bacteria and *S. sonnei*.

In conclusion, multiple XDR clones of *S. sonnei* have emerged and are co-circulating in Vietnam. Commensal *E. coli* in the gastrointestinal tract of Vietnamese children display an exceptionally high degree of diversity in AMR genes and plasmid composition, and our evidence suggests these are the most likely reservoir for the maintenance and transfer of MDR plasmids to cipR *S. sonnei*. Our data further suggest *in vivo* plasmid transfer between commensal *E. coli* and cipR *S. sonnei* during infection, which is likely facilitated by the presence of sub-MIC concentrations of ciprofloxacin. We advocate for the continued surveillance of XDR *S. sonnei* in Vietnam and a suggest a urgent re-evaluation of the empirical use of ciprofloxacin for a range of gastrointestinal infections.

## Acknowledgements

This work was funded by Sir Henry Dale Fellow, jointly funded by the Wellcome Trust and the Royal Society, to SB (100087/Z/12/Z) and an Oak Leader Fellowship to DPT.

## Author contributions

PTD, MAR and SB designed the study. PTD performed in data analysis and interpretation of the results under the scientific guidance of MAR and SB. PTD drafted and edited the paper, with MAR and SB revising and structuring the paper. TNTN, HCT, CB, FA, HNDT, and HTT performed laboratory work and generated the data for analysis. DVT recruited patients and performed the clinical work required for the study. HCT, CB, and GET contributed to the editing of the paper. All authors read and approved the final draft.

## Additional information

None

---

**Supplementary Figure 1.**
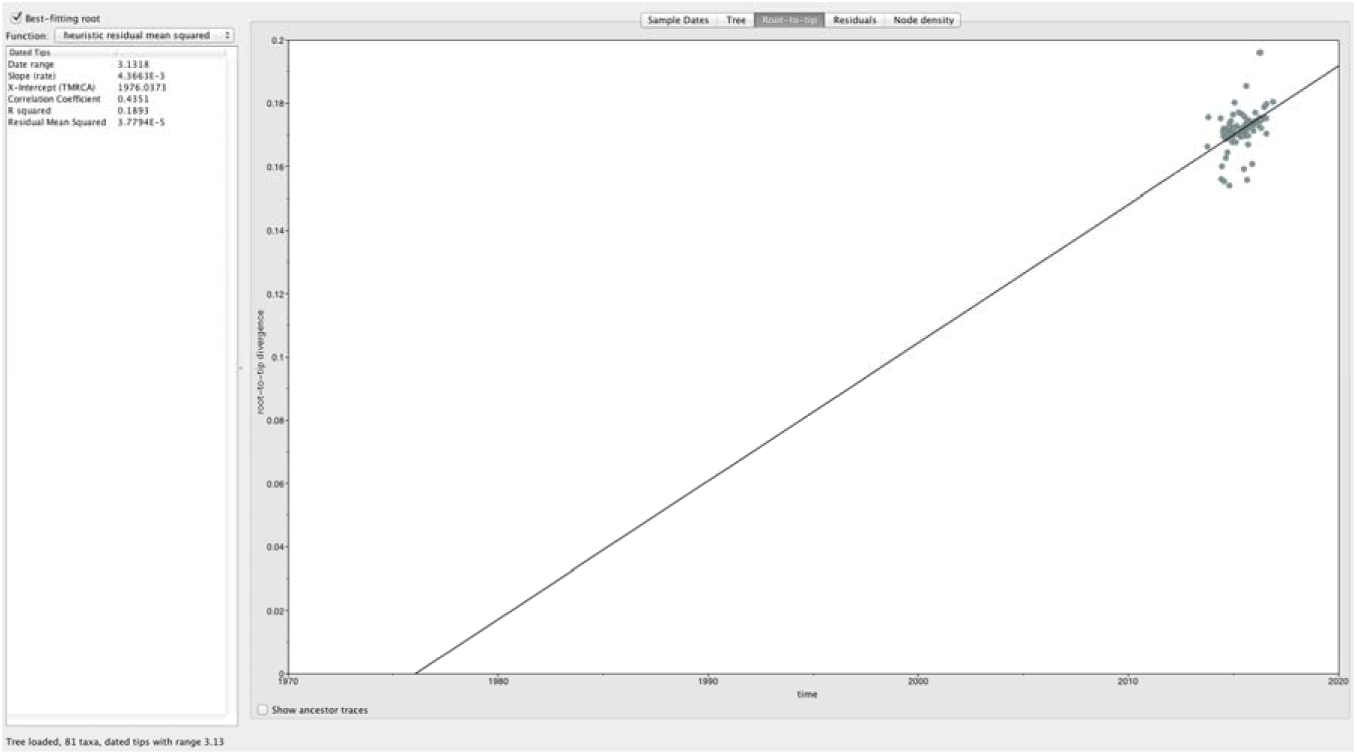
Root-to-tip regression for the maximum likelihood tree of *Shigella sonnei* in Vietnam, 2014 and 2016 Each point on the plot corresponds to a measurement of genetic distance from the inferred root to each tip in the tree. The solid line is the regression line fitted using the ordinary least squares method. The slope of the line is a crude estimate of the evolutionary rate, the x-intercept corresponds to the time to the most recent common ancestor, and the R^2^ value measures the degree of clock-like behavior.

**Supplementary Figure 2.**
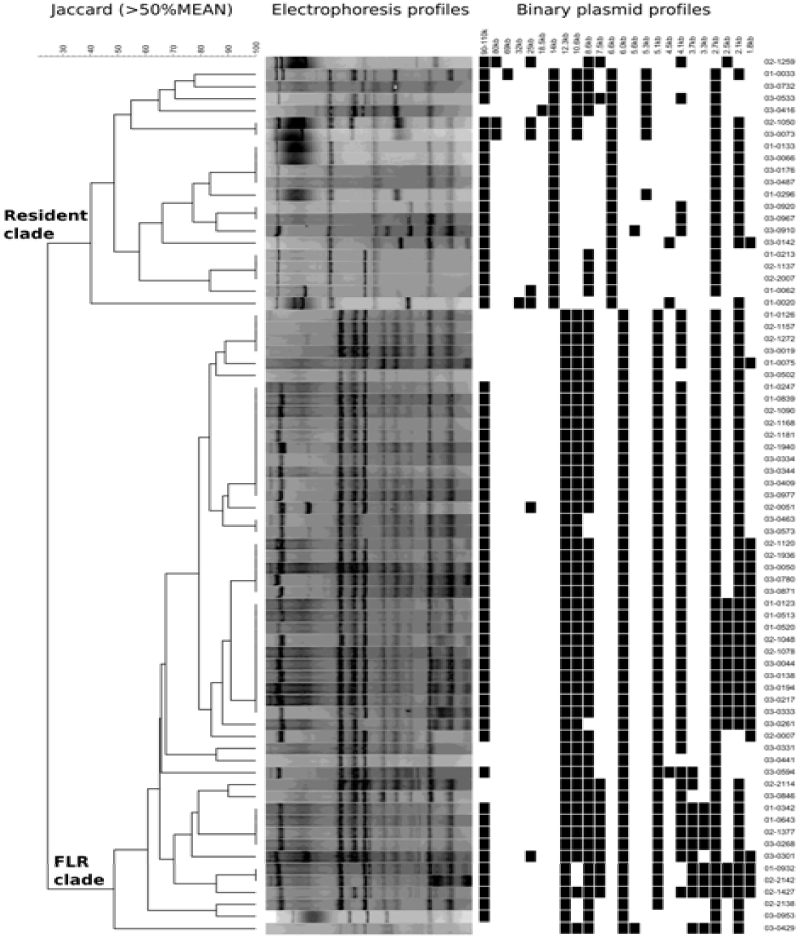
Plasmid profiling of *Shigella sonnei* in Vietnam, 2014 and 2016 Dendrogram shows the difference in plasmid electrophoresis patterns between the resident *S. sonnei* clade and the cipR *S. sonnei* clade in Vietnam. Cluster analysis was performed with Bionumerics by using the Jaccard coefficient and the unweighted pair group mathematical average (UPGMA) clustering algorithm.

**Supplementary Figure 3.**
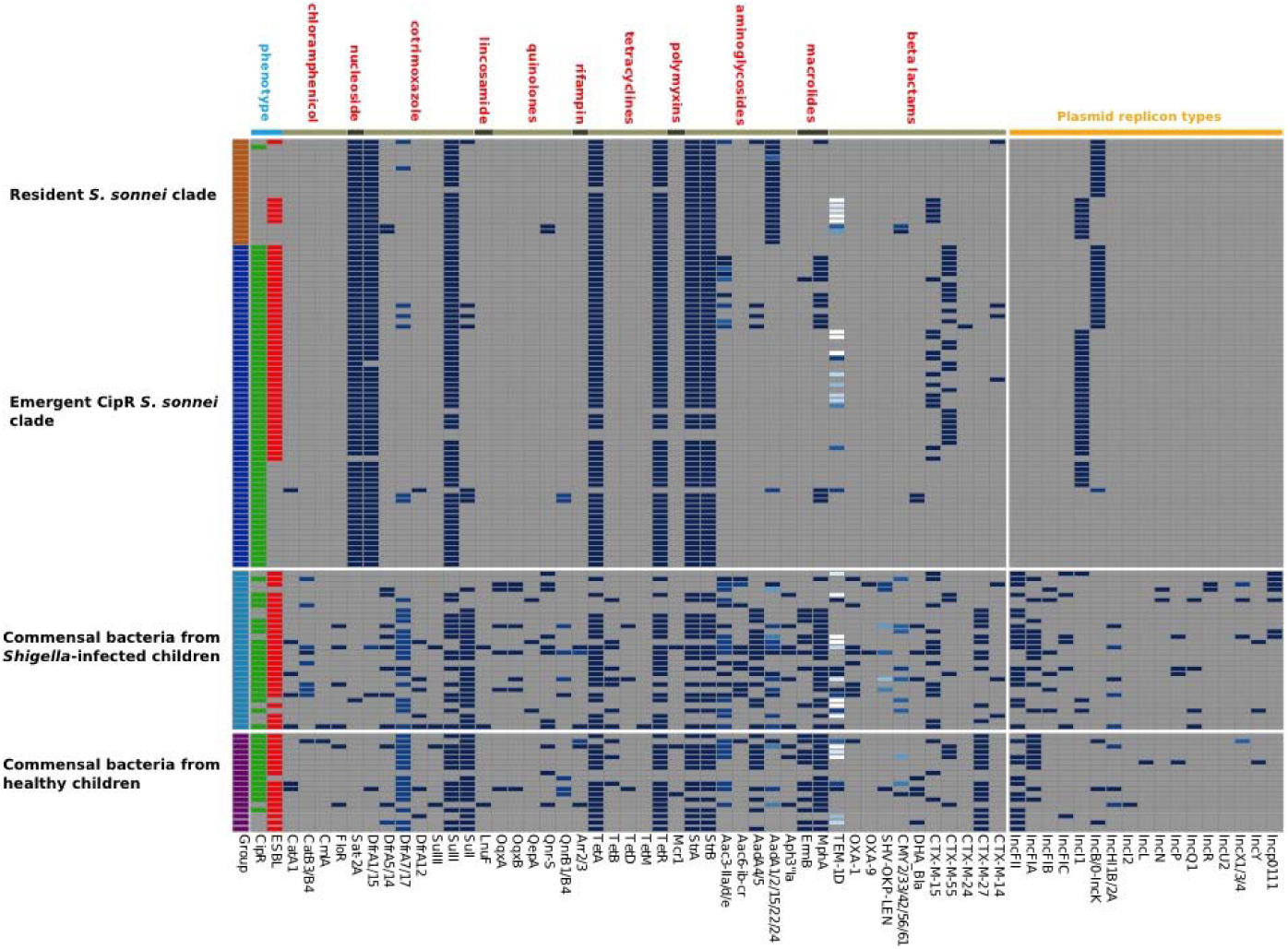
Distribution of antimicrobial resistance genes and plasmid groups in *Shigella sonnei* and human commensal bacteria The first column highlights the four different isolate types in different colors. Fecal/rectal swab cultures with ciprofloxacin-resistant and ESBL-producing isolates are highlighted in green and red (second and third columns, respectively). The remaining columns show the distribution of all antimicrobial resistance genes and plasmid groups identified in commensal bacteria and *S. sonnei*. AMR genes are grouped together based on the class of antimicrobial agents to which they are resistant, with different variants of an antimicrobial resistant gene shown in different shades of blue.

## References

1 Kotloff KL, Winickoff JP, Ivanoff B, et al. Global burden of Shigella infections: Implications for vaccine development and implementation of control strategies. Bull World Health Organ 1999; 77: 651–66.

2 Lanata CF, Fischer-Walker CL, Olascoaga AC, et al. Global causes of diarrheal disease mortality in children. PLoS One 2013; 8: e72788.

3 Kotloff KL, Nataro JP, Blackwelder WC, et al. Burden and aetiology of diarrhoeal disease in infants and young children in developing countries (the Global Enteric Multicenter Study, GEMS): a prospective, case-control study. Lancet 2013; 382: 209–22.

4 Thompson CN, Duy PT, Baker S. The rising dominance of Shigella sonnei: An intercontinental shift in the etiology of bacillary dysentery. PLoS Negl. Trop. Dis. 2015; 9. DOI:10.1371/journal.pntd.0003708.

5 Baker KS, Campos J, Pichel M, et al. Whole genome sequencing of *Shigella sonnei* through PulseNet Latin America and Caribbean: advancing global surveillance of foodborne illnesses. Clin Microbiol Infect 2017; 23: 845–53.

6 Holt KE, Baker S, Weill F-X, et al. Shigella sonnei genome sequencing and phylogenetic analysis indicate recent global dissemination from Europe. 2012; 44. DOI:10.1038/ng.2369.

7 D. Legros, D. Legros. Guidelines for the control of shigellosis, including epidemics due to Shigella dysenteriae type 1. World Health 2005; : 1–70.

8 Pazhani GP, Ramamurthy T, Mitra U, Bhattacharya SK, Niyogi SK. Species diversity and antimicrobial resistance of Shigella spp. isolated between 2001 and 2004 from hospitalized children with diarrhoea in Kolkata (Calcutta), India. Epidemiol Infect 2005; 133: 1089–95.

9 Shakya G, Acharya J, Adhikari S, Rijal N. Shigellosis in Nepal: 13 years review of nationwide surveillance. J Health Popul Nutr 2016; 35: 36.

10 Von Seidlein L, Deok RK, Ali M, et al. A multicentre study of Shigella diarrhoea in six Asian countries: Disease burden, clinical manifestations, and microbiology. PLoS Med 2006; 3: 1556–69.

11 Pazhani GP, Niyogi SK, Singh AK, et al. Molecular characterization of multidrug-resistant Shigella species isolated from epidemic and endemic cases of shigellosis in India. J Med Microbiol 2008; 57: 856–63.

12 Rajpara N, Nair M, Chowdhury G, et al. Molecular analysis of multidrug resistance in clinical isolates of *Shigella* spp. from 2001-2010 in Kolkata, India: role of integrons, plasmids, and topoisomerase mutations. Infect Drug Resist 2018; Volume 11: 87–102.

13 De Lappe N, O’Connor J, Garvey P, McKeown P, Cormican M. Ciprofloxacin-resistant Shigella sonnei associated with travel to India. Emerg Infect Dis 2015; 21: 894–6.

14 Bowen A, Hurd J, Hoover C, et al. Importation and Domestic Transmission of Shigella sonnei Resistant to Ciprofloxacin – United States, May 2014-February 2015. MMWR Morb Mortal Wkly Rep 2015; 64: 318–20.

15 Nüesch-Inderbinen M, Heini N, Zurfluh K, Althaus D, Hächler H, Stephan R. Shigella antimicrobial drug resistance mechanisms, 2004–2014. Emerg Infect Dis 2016; 22: 1083–5.

16 Ruekit S, Wangchuk S, Dorji T, et al. Molecular characterization and PCR-based replicon typing of multidrug resistant Shigella sonnei isolates from an outbreak in Thimphu, Bhutan. BMC Res Notes 2014; 7. DOI:10.1186/1756-0500-7-95.

17 Kim JS, Kim JJ, Kim SJ, et al. Outbreak of ciprofloxacin-resistant Shigella sonnei associated with travel to Vietnam, Republic of Korea. Emerg Infect Dis 2015; 21: 1247–50.

18 Gaudreau C, Ratnayake R, Pilon PA, Gagnon S, Roger M, Levesque S. Ciprofloxacin-resistant Shigella sonnei among men who have sex with men, Canada, 2010. Emerg Infect Dis 2011; 17: 1747–50

19 Chiou CS, Izumiya H, Kawamura M, et al. The worldwide spread of ciprofloxacin-resistant Shigella sonnei among HIV-infected men who have sex with men, Taiwan. Clin Microbiol Infect 2016; 22: 383.e11–383.e16.

20 Baker KS, Dallman TJ, Field N, et al. Genomic epidemiology of Shigella in the United Kingdom shows transmission of pathogen sublineages and determinants of antimicrobial resistance. Sci Rep 2018; 8: 1–8.

21 Bodhidatta L, Thanh TH, Thanh DP, et al. Introduction and establishment of fluoroquinolone-resistant Shigella sonnei into Bhutan. Microb Genomics 2015; 1. DOI:10.1099/mgen.0.000042.

22 Chung The H, Rabaa MA, Pham Thanh D, et al. South Asia as a Reservoir for the Global Spread of Ciprofloxacin-Resistant Shigella sonnei: A Cross-Sectional Study. PLoS Med 2016; 13: 1–12.

23 Duong VT, Tuyen HT, Van Minh P, et al. No Clinical benefit of empirical antimicrobial therapy for pediatric diarrhea in a high-usage, high-resistance setting. Clin Infect Dis 2018; 66: 504–11.

24 Magiorakos AP, Srinivasan A, Carey RB, et al. Multidrug-resistant, extensively drug-resistant and pandrug-resistant bacteria: An international expert proposal for interim standard definitions for acquired resistance. Clin Microbiol Infect 2012; 18: 268–81.

25 Thiem VD, Sethabutr O, Von Seidlein L, et al. Detection of Shigella by a PCR Assay Targeting the ipaH Gene Suggests Increased Prevalence of Shigellosis in Nha Trang, Vietnam. J Clin Microbiol 2004; 42: 2031–5.

26 Thompson CN, Anders KL, Nhi LTQ, et al. A cohort study to define the age-specific incidence and risk factors of Shigella diarrhoeal infections in Vietnamese children: A study protocol. BMC Public Health 2014; 14. DOI:10.1186/1471-2458-14-1289.

27 Li H, Handsaker B, Wysoker A, et al. The Sequence Alignment/Map format and SAMtools. Bioinformatics 2009; 25: 2078–9.

28 Croucher NJ, Page AJ, Connor TR, et al. Rapid phylogenetic analysis of large samples of recombinant bacterial whole genome sequences using Gubbins. Nucleic Acids Res 2015; 43: e15.

29 Nguyen LT, Schmidt HA, Von Haeseler A, Minh BQ. IQ-TREE: A fast and effective stochastic algorithm for estimating maximum-likelihood phylogenies. Mol Biol Evol 2015; 32: 268–74.

30 Rambaut A, Lam TT, Max Carvalho L, Pybus OG. Exploring the temporal structure of heterochronous sequences using TempEst (formerly Path-O-Gen). Virus Evol 2016; 2: vew007.

31 Drummond AJ, Rambaut A. BEAST: Bayesian evolutionary analysis by sampling trees. BMC Evol Biol 2007; 7. DOI:10.1186/1471-2148-7-214.

32 Drummond AJ, Rambaut A, Shapiro B, Pybus OG. Bayesian Coalescent Inference of Past Population Dynamics from Molecular Sequences. DOI:10.1093/molbev/msi103.

33 Drummond AJ, Ho SYW, Phillips MJ, Rambaut A. Relaxed Phylogenetics and Dating with Confidence. DOI:10.1371/journal.pbio.0040088.

34 Xie W, Lewis PO, Fan Y, Kuo L, Chen MH. Improving marginal likelihood estimation for bayesian phylogenetic model selection. Syst Biol 2011; 60: 150–60.

35 Lartillot N, Philippe H. Computing Bayes factors using thermodynamic integration. Syst Biol 2006; 55: 195–207.

36 Inouye M, Dashnow H, Raven L-A, et al. SRST2: Rapid genomic surveillance for public health and hospital microbiology labs. Genome Med 2014; 6: 90.

37 Gupta SK, Padmanabhan BR, Diene SM, et al. ARG-annot, a new bioinformatic tool to discover antibiotic resistance genes in bacterial genomes. Antimicrob Agents Chemother 2014; 58: 212–20.

38 Carattoli A, Zankari E, Garciá-Fernández A, et al. In Silico detection and typing of plasmids using plasmidfinder and plasmid multilocus sequence typing. Antimicrob Agents Chemother 2014; 58: 3895–903.

39 García-Fernández A, Chiaretto G, Bertini A, et al. Multilocus sequence typing of IncI1 plasmids carrying extended-spectrum β-lactamases in Escherichia coli and Salmonella of human and animal origin. J Antimicrob Chemother 2008; 61: 1229–33.

40 Zerbino DR, Birney E. Velvet: Algorithms for de novo short read assembly using de Bruijn graphs. Genome Res 2008; 18: 821–9.

41 Seemann T. Prokka: Rapid prokaryotic genome annotation. Bioinformatics 2014; 30: 2068–9.

42 Assefa S, Keane TM, Otto TD, Newbold C, Berriman M. ABACAS: Algorithm-based automatic contiguation of assembled sequences. Bioinformatics 2009; 25: 1968–9.

43 Carver T, Berriman M, Tivey A, et al. Artemis and ACT: Viewing, annotating and comparing sequences stored in a relational database. Bioinformatics 2008; 24: 2672–6.

44 Wood DE, Salzberg SL. Kraken: Ultrafast metagenomic sequence classification using exact alignments. Genome Biol 2014; 15. DOI:10.1186/gb-2014-15-3-r46.

45 CI Kado and ST. Liu. Rapid Procedure for Detection and Isolation of Large and Small Plasmids. J Bacteriol 1981; 145: 1365–73.

46 Bankevich A, Nurk S, Antipov D, et al. SPAdes: A New Genome Assembly Algorithm and Its Applications to Single-Cell Sequencing. J Comput Biol 2012; 19: 455–77.

47 Holt KE, Thieu Nga TV, Thanh DP, et al. Tracking the establishment of local endemic populations of an emergent enteric pathogen. Proc Natl Acad Sci U S A 2013; 110: 17522–7.

48 Nhi LTQ, Tuyen HT, Trung PD, et al. Excess body weight and age associated with the carriage of fluoroquinolone and third-generation cephalosporin resistance genes in commensal escherichia coli from a cohort of urban vietnamese children. J Med Microbiol 2018; 67: 1457–66.

49 Bearson BL, Brunelle BW. Fluoroquinolone induction of phage-mediated gene transfer in multidrug-resistant Salmonella. Int J Antimicrob Agents 2015; 46: 201–4.

50 Hastings PJ, Rosenberg SM, Slack A. Antibiotic-induced lateral transfer of antibiotic resistance. Trends Microbiol. 2004; 12: 401–4.

51 Qin T, Kang H, Ma P, Li P, Huang L, Gu B. SOS response and its regulation on the fluoroquinolone resistance. Ann Transl Med 2015; 3: 358.

52 Baharoglu Z, Bikard D, Mazel D. Conjugative DNA transfer induces the bacterial SOS response and promotes antibiotic resistance development through integron activation. PLoS Genet 2010; 6: 1–10.

53 Ubeda C, Maiques E, Knecht E, Lasa I, Novick RP, Penadés JR. Antibiotic-induced SOS response promotes horizontal dissemination of pathogenicity island-encoded virulence factors in staphylococci. Mol Microbiol 2005; 56: 836–44.

54 Beaber JW, Hochhut B, Waldor MK. SOS response promotes horizontal dissemination of antibiotic resistance genes. Nature 2004; 427: 72–4.

55 Dalhoff A. Global fluoroquinolone resistance epidemiology and implictions for clinical use. Interdiscip Perspect Infect Dis 2012; 2012. DOI:10.1155/2012/976273

56 Thanh DP, Karkey A, Dongol S, et al. A novel ciprofloxacin-resistant subclade of h58. Salmonella typhi is associated with fluoroquinolone treatment failure. Elife 2016; 5. DOI:10.7554/eLife.14003.

57 Talukder KA, Khajanchi BK, Islam MA, et al. Genetic relatedness of ciprofloxacin-resistant Shigella dysenteriae type 1 strains isolated in south Asia. J Antimicrob Chemother 2004; 54: 730–4.

58 Holden MTG, Hsu L-Y, Kurt K, et al. A genomic portrait of the emergence, evolution, and global spread of a methicillin-resistant Staphylococcus aureus pandemic. Genome Res 2013; 23: 653–64.

59 Banerjee R, Johnson JR. A new clone sweeps clean: The enigmatic emergence of Escherichia coli sequence type 131. Antimicrob. Agents Chemother. 2014; 58: 4997–5004.

60 Le Hello S, Bekhit A, Granier SA, et al. The global establishment of a highly-fluoroquinolone resistant Salmonella enterica serotype Kentucky ST198 strain. Front Microbiol 2013; 4. DOI:10.3389/fmicb.2013.00395.

61 He M, Miyajima F, Roberts P, et al. Emergence and global spread of epidemic healthcare-associated Clostridium difficile. Nat Genet 2013; 45: 109–13

62 Xia L, Liu Y, Xia S, et al. Prevalence of ST1193 clone and IncI1/ST16 plasmid in E-coli isolates carrying blaCTX-M-55gene from urinary tract infections patients in China. Sci Rep 2017; 7. DOI:10.1038/srep44866.

63 Kim Y, Oh T, Nam YS, Cho SY, Lee HJ. Prevalence of ST131 and ST1193 among bloodstream isolates of Escherichia coli not susceptible to ciprofloxacin in a tertiary care university hospital in korea, 2013-2014. Clin Lab 2017; 63: 1541–3.

64 Wu J, Lan F, Lu Y, He Q, Li B. Molecular characteristics of ST1193 clone among phylogenetic group B2 Non-ST131 fluoroquinolone-resistant Escherichia coli. Front Microbiol 2017; 8. DOI:10.3389/fmicb.2017.02294.

65 Tchesnokova VL, Rechkina E, Larson L, et al. Rapid and Extensive Expansion in the U.S. of a New Multidrug-Resistant Escherichia coli Clonal Group, Sequence Type ST1193. Clin Infect Dis 2018; : 2016–9.

66 Zurita J, Ortega-Paredes D, Barba P. First description of *Shigella sonnei* harboring blaCTX-M-55 outside Asia. J Microbiol Biotechnol 2016; 26: 2224–7.

67 Lee W, Chung HS, Lee H, et al. CTX-M-55-type extended-spectrum β-lactamase-producing Shigella sonnei isolated from a Korean patient who had travelled to China. Ann Lab Med 2013; 33: 141–4.

68 Qu F, Ying Z, Zhang C, et al. Plasmid-encoding extended-spectrum β-lactamase CTX-M-55 in a clinical Shigella sonnei strain, China. Future Microbiol 2014; 9: 1143–50.

69 Praszkier J, Pittard AJ. Control of replication in I-complex plasmids. Plasmid. 2005; 53: 97–112.

70 Allué-Guardia A, Koenig SSK, Quirós P, Muniesa M, Bono JL, Eppinger M. Closed genome and comparative phylogenetic analysis of the clinical multidrug resistant Shigella sonnei strain 866. Genome Biol Evol 2018. DOI:10.1093/gbe/evy168.

71 Folster JP, Pecic G, Krueger A, et al. Identification and characterization of CTX-M-producing Shigella isolates in the United States. Antimicrob. Agents Chemother. 2010; 54: 2269–70.

72 Ma Q, Xu X, Luo M, et al. A waterborne outbreak of shigella sonnei with resistance to azithromycin and third-generation cephalosporins in China in 2015. Antimicrob Agents Chemother 2017; 61. DOI:10.1128/AAC.00308-17.

73 Seral C, Rojo-Bezares B, Garrido A, Gude MJ, Sáenz Y, Castillo FJ. Characterisation of a CTX-M-15-producing Shigella sonnei in a Spanish patient who had not travelled abroad. Enferm Infecc Microbiol Clin 2012; : 2011–3

74 Kim JS, Kim J, Jeon SE, et al. Complete nucleotide sequence of the IncI1 plasmid pSH4469 encoding CTX-M-15 extended-spectrum β-lactamase in a clinical isolate of Shigella sonnei from an outbreak in the Republic of Korea. Int J Antimicrob Agents 2014; 44: 533–7.

75 Johnson TJ, Wannemuehler YM, Johnson SJ, et al. Plasmid replicon typing of commensal and pathogenic Escherichia coli isolates. Appl Environ Microbiol 2007; 73: 1976–83.

76 Ravi A. Characterization of the infant gut microbiota mobilome. 2017. http://hdl.handle.net/11250/2497973.

77 Bratoeva MP, John JF. In vivo R–plasmid transfer in a patient with a mixed infection of shigella dysentery. Epidemiol Infect 1994; 112: 247–52.

78 Rashid H, Rahman M. Possible transfer of plasmid mediated third generation cephalosporin resistance between Escherichia coli and Shigella sonnei in the human gut. Infect Genet Evol 2015; 30: 15–8.

79 Duong VT, Tuyen HT, Van Minh P, et al. No Clinical benefit of empirical antimicrobial therapy for pediatric diarrhea in a high-usage, high-resistance setting. Clin Infect Dis 2018; 66: 504–11.

